# Molecular surveillance of HIV, HBV, and HCV amongst blood donors in five Chinese regions

**DOI:** 10.1101/2020.01.22.916320

**Authors:** Xiaoting Lv, Mary A Rodgers, Peng Yin, Ling Ke, Ping Fu, Binting Wu, Yu Liu

## Abstract

Hepatitis B (HBV), hepatitis C (HCV) and human immunodeficiency virus (HIV) are transfusion transmissible infections (TTIs) agents that threaten the safety of the blood supply. Surveillance of the variance of those viruses is an important way to monitor their diversity and evolution to improve safety in blood transfusion. In this study, we characterized the specimens of blood donors from 13 blood centers located in 5 Chinese regions.

Samples collected between 2014 and 2017 were screened with serological and molecular tests conducted on Abbott ARCHITECT and m2000 platforms. Sequencing was used to determine the classifications. The HBV immune escape mutations were also analyzed for assessing vaccine breakthrough risks and challenges for diagnostic tests. For HIV, 11 genotypes or recombinants were identified. The predominant genotype was C, which accounts for 42%. For HBV, the genotypes of B, C and D were identified, with B and C predominating. The major subgenotype was B2, comprising 84.1% of all infections. 79 out of 113 (69.9%) samples carried escape mutations in the “a” determinant region with 69 (87.3%) multiple mutants and 15 (19%) escape mutants which will affect HBsAg detection. For HCV, 7 genotypes or subtypes were identified. The major genotype was 1b (48%), followed by 6a (16.7%) and 2a & 3a (10%). This study provides the information of diversity of HBV, HCV and HIV strains circulating in blood centers from 5 regions in China. These data can also be scientific basis for development of detection assays that mitigate the impact of viral diversity on performance.

**Importance:** The prevalence of TTIs in blood donations is important for evaluating blood safety and it can also reflect the burden of these disease among populations. Virus variance is threat to blood safety due to it may affect assays detection by nucleic acid, antigen and antibody-based methods in blood donors. HIV, HBV and HCV exhibit high degrees of genetic diversity, with different strains predominating in different geographic locations. The aim of this study is to assess the diversity of HBV, HCV and HIV among blood donors in China. In this study, 13 blood centers located in 5 Chinese regions were involved and the most informative phylogenetic regions of each virus had been sequenced. This will benefit for viral monitoring by subtype/genotype analyses to determine whether the distributions of variants are changing over time and geographically, and to speculate whether previously rare subtypes are becoming established in blood donors in China.

## Introduction

Hepatitis B (HBV), hepatitis C (HCV) and human immunodeficiency virus (HIV) are three important transfusion-transmitted infectious agents which are threats to blood safety in China (1). Before 1998, a high number of HBV, HCV, and HIV infections in China were from a blood transfusion. Since the three agents were included for blood screening, especially nucleic acid testing of the three agents conducted for all blood donors, the infection of the three agents through blood transfusion decreased significantly. However, the threat of these agents to blood safety still exists in China (2).

Several studies have reported the molecular epidemiological data of the three agents in general populations in China. The first nationwide molecular epidemiological survey of HIV conducted in 1996-1998 showed that subtype B’/B (47.5%), subtype C (34.3%), and CRF01_AE (9.6%) were the most predominant strains in China (3). However, the second and third surveys conducted in 2002-2003 and 2006 indicated that the proportions of HIV-1 genotypes had sharply shifted. The subtype B’/B and C have been replaced by CRF07_BC, CRF01_AE, and CRF08_BC (4, 5).

HBV and HCV genotypes have geographic distribution characteristics (6-8). HBV has 9 genotypes (A-I) (9) and more than 35 subgenotypes that have been identified (10). Genotypes B and C are the predominant HBV strains in China. North of the Qinling Mountains-Huaihe River Line, genotype C (75.3%) is dominating with a small portion of types B (23.4%) and D (1.3%) (11). In the south of China, the major genotype is B (12). The distribution of HCV genotypes is closely related to the modes of HCV acquisition (13). In the late 1980s and early 1990s in China, illegal blood collection and donation greatly promoted the spread of HCV genotype 1 and 2 (14). In 1998, mandatory HCV screening of blood and blood products was implemented in China, gradually changing the predominant mode of HCV transmission from direct contact with human blood to intravenous and percutaneous drugs used (15). Consequently, the HCV genotype distribution in China has been shifting over time, especially in the southwestern provinces (16). In China, the only multicenter investigation of the distribution of HCV genotypes, conducted in 2011, indicated that the most prevalent HCV subtype was 1b, followed by 2a (17). Over time, the prevalence of genotypes 3, 6, and new mixed-infection genotype has also been increasing (18).

These data indicate the diversity of HIV, HBV, and HCV and the importance of monitoring virus diversity and evolution for blood safety in China. Except for the diversity of genotypes, mutations in the sequences of these agents may lead to false-negative detection in blood screening and introduce TTIs that jeopardizes the blood safety. Take the mutants of HBV as an example, the mutants in “a” determinant region which located in the major immunogenic region can lead to false-negative detection, vaccine escape and may evade natural immune responses (19). In previous reports, the sG145R mutant is the most widely reported. Besides sG145R, the K141E, T131I, and insertion of amino acids between 123 and 124 can affect the structure of HBsAg to cause vaccine escape as well (20). Recently, other mutants in “a”-determinant region are linked to vaccine escape, like T116N, P120S/E, I/T126A/N/I/T126A/N/I/S, Q129H/R, M133L, K141E, P142S, and D144A/E (21)

Considering the importance of the molecular epidemiological characteristics of these agents in blood donors for blood safety, we performed molecular surveillance of HIV, HBV, and HCV to determine the genotype diversity and characteristics of HBV S region amongst blood donors in 5 Chinese regions.

## Methods

### Samples

Retrospective leftover plasma from 216 HIV, 207 HBV, or 204 HCV available positive blood donations collected from 13 blood centers in four provinces and one municipality (Chongqing, Guangxi, Henan, Sichuan, Xinjiang) between 2014 and 2017 were included in the study. All positive donations were screened with two ELISA assays as previously described at the blood centers where they were first collected (22). These samples were confirmed positive in the laboratory of IBT. The viral load and serology screening tests were performed on Abbott ARCHITECT (HIV Combo Ag/Ab, HBsAg, and anti-HCV) and m2000 (RealTime HIV-1, HBV, HCV) instruments per manufacturer’s instructions. This study was approved by the IRB of the Institute of Blood Transfusion. The written consent was obtained from all donors at the time of donation.

### Viral nucleic acid extraction and amplification

HBV DNA was extracted from 200 μL plasma using a QIA amp ^®^ DNA Blood Mini Kit (QIAGEN, Hilden, Germany) according to the manufacturer’s instructions and 50 μL of eluted DNA was stored at −70 °C until use. The HBV S and Pol regions were amplified by PCR using a thermal cycler (Veriti, Applied Biosystems, MA, USA) with 56F-CCTGCTGGTGGCTCCAGTTC, 645F-TATGTTGCCCGTTTGTCCTCTAAT as the forward primer and 1253R-GCAGTATGGATCGGCAGAGGAG,1580R-AGGTGAAGCGAAGTGCACACG as the reverse primer respectively for the first round PCR. The first round PCR was performed in a total volume of 50 μL, with the following reaction: denaturation at 95 °C for 10 minutes, followed by 35 cycles of 15 seconds of denaturation at 95 °C, 30 seconds of annealing at 53 °C, and 30 seconds of extension at 72 °C, with a final extension at 72 °C for 5 minutes. The second round PCR condition is the same as the first round PCR. The forward primers is 178F-CCTAGGACCCCTGCTCGTGTTACAGGC, 784F-TCCCTTTATACCGCTGTTACCAAT and reverse primer are 1186R-CCAGTGGGGGTTGCRTCAGC, 1421R-CGCYGACGGGACGTARACAA in S and Pol region, respectively.

HCV and HIV RNA were extracted from 200 μL plasma using a QIA amp^®^ Viral RNA Mini Kit (QIAGEN, Hilden, Germany) according to the manufacture’s instruction and 50 μL of eluted RNA was stored at −70 °C until use. The HCV 5’UTR-core was amplified, one-step RT-PCR was performed with 15 μL of RNA, forward primer 127F-TCCCGGGAGAGCCATAGT, reverse primers 852Rb-AGGAAGATAGAGAAAGAGCAACC and 852Rc-AGGAAGATAGAAAAGGAGCAACC using QIAGEN One-Step RT-PCR kit (QIAGEN GmbH, Hilden, Germany) according to the manufacturer’s instructions. Cycling conditions were 50 °C for 30 minutes, 95 °C for 15 minutes, 50 cycles of 94 °C for 15 seconds, 50 °C for 30 seconds and 72 °C for 1 minute 30 seconds, and final extension at 72 °C for 10 minutes.

The env IDR (immunodominant region of gp41), pol integrase and 5’LTR were amplified with forwarding primers JH35F-TGARGGACAATTGGAGAARTGA, Poli5-CACACAAAGGTATTGGAGGAAATG, MGAGrev-GCTCTCGCACCCATCTCTCT respectively; and reverse primers JH38R-GGTGARTATCCCTKCCTAAC, Poli8-TAGTGGGATGTGTACTTCTGAAC and MRU3fwd-GAGCCTGGGAGCTCTCTG. The cycling conditions for env IDR are 50 °C for 30 minutes, then 95 °C for 15 minutes and 50 cycles with 94 °C for 15 seconds, 50 °C for 30 seconds and 72 °C for 1 minute. Finalize with 72 °C for 7 minutes. The condition for Pol Integrase is 50 °C for 30 minutes, then 95 °C for 15 minutes and 50 cycles with 94 °C for 15 seconds, 60 °C for 30 seconds and 72 °C for 1 minute. Finalize with 72 °C for 7 minutes. 5UTR cycling condition is 50 °C for 30 minutes, then 95 °C for 15 minutes and 50 cycles with 94 °C for 15 seconds, 55 °C for 30 seconds and 72 °C for 1 minute. Finalize with 72 °C for 7 minutes.

### Sequencing

Nucleotide fragments amplified from above were tested for molecular characterization by Sanger sequencing of subgenomic amplicons by Sangon Biotech (Shanghai, China).

### HIV phylogenetic classification

Viral sequences were aligned with reference strains, including all subtypes and CRFs 1-90 by CLUSTALW. Alignments were manually edited in BioEdit 7.0.4.1 or higher to remove gaps and Neighbor-Joining phylogenetic inference was performed using PHYLIP 3.5c (J. Felsenstein, University of Washington, Seattle, USA). Evolutionary distances were estimated with Dnadist (Kimura two-parameter method) and phylogenetic relationships were determined by Neighbor (neighbor-joining method). The Branch reproducibility of trees was evaluated using Seqboot (100 replicates) and Consense. Programs were run with default parameters. For specimens clustering outside of the CRF02_AG reference branch, a second alignment and phylogenetic tree analysis was completed, including all subtypes and 72 CRFs. Classification was assigned for specimens with bootstrap support of 70 or higher. Phylogenetic trees were visualized in FigTree v1.4.2 (University of Edinburgh, UK).

### HCV phylogenetic classification

Sequence data were edited and assembled using Sequencher version 5.4.6 (Gene Codes Corp, Ann Arbor, MI, USA). Sequences were aligned to N=117 reference strains, including subgenotypes of HCV genotypes 1-7 by MUSCLE in Sequencher. Gaps were manually removed in BioEdit version 7.0.4.1(25), and Neighbor-Joining phylogenetic inference was performed using the PHYLIP 3.5c (J. Felsenstein, University of Washington, Seattle, WA, USA) software package as previously described (23).

### HBV phylogenetic classification and mutation analysis

Sequence data was analyzed using Sequencher version 5.2.3 or higher (Gene Codes Corp., Ann Arbor, MI). To determine the genotype of HBV, the DNA sequences were aligned with genotype reference sequences using the CLUSTALW method and manually edited in BioEdit 7.0.4.1 or higher (24). Phylogenetic trees were prepared as described above for HIV and visualized in FigTree v1.4.2 (University of Edinburgh, UK). HBsAg subtype was determined as described in a previous paper (25). HBsAg escape mutations were identified using the web-based program Geno2pheno (hbv) v2.0 (Informatik).

## Results

### Demographics

To determine the HIV, HBV, and HCV strains present amongst blood donors in China, seropositive plasma specimens were collected from 13 blood centers in Chongqing, Guangxi, Henan, Sichuan, and Xinjiang regions between 2014 and 2017.

Specimens were identified as HIV antibody, HBV surface antigen (HBsAg), or HCV antibody positive at each collection site by two independent ELISA tests. Subsequent viral load testing with the m2000 RealTi*m*e HIV, HBV, or HCV test (Abbott Molecular Diagnostics, Des Plaines, IL, USA) confirmed viral nucleic acids were present in a total of 626 specimens (N=215 HIV, N=207 HBV, N=204 HCV) for further characterization of the viral strains present. Subgenomic direct amplicon sequencing was performed for specimens with sufficient viral load, and at least one sequence was obtained for N=107 HIV, N=113 HBV, and N=60 HCV specimens (Table 1). The collection sites and demographics of the donors of these sequenced specimens were summarized in Figure 1. The majority of donors were male (N=200, 71.4%), Han (N=257, 91.8%), and from Chongqing (N=179, 63.9%). Donor ages ranged from 18 to 59 years old, and the average age was 35 Junior, Bachelor and High school students account for 235 (84%). Technical secondary school, Primary school, Master and Bachelor Junior account for 26 (9%), Other 19 (6.79%) donors were unknown education status. The most represented occupation was freelancers 72 (25.71%), followed by students 42 (15%), Agriculture, Fisheries and Forestry 30 (10.71%), and Commercial service 30 (10.71%).

**Table 1.** Blood donors from 13 blood centers of 4 provinces and 1 municipality

| Provinces/Municipality | Characteristics |  |  | Grand Total |
| --- | --- | --- | --- | --- |
|  | HBV | HCV | HIV |  |
| **Chongqing** | **112** | **26** | **41** | **179** |
| Chongqing Blood Center | 112 | 26 | 41 | 179 |
| **Guangxi** |  | **20** | **9** | **29** |
| Liuzhou Blood Center |  | 20 | 9 | 29 |
| **Henan** |  | **2** | **18** | **20** |
| Henan Red Cross Blood Center |  |  | 15 | 15 |
| Luoyang Blood Center |  | 2 | 3 | 5 |
| **Sichuan** | **1** | **10** | **34** | **45** |
| Bazhong Blood Center |  |  | 3 | 3 |
| Chengdu Blood Center |  |  | 6 | 6 |
| Deyang Blood Center |  |  | 5 | 5 |
| Guangyuan Blood Center |  |  | 4 | 4 |
| Meishan Blood Center |  |  | 7 | 7 |
| Mianyang Blood Center | 1 | 10 | 4 | 15 |
| Nanchong Blood Center |  |  | 1 | 1 |
| Ya'an Blood Center |  |  | 4 | 4 |
| **Xinjiang** |  | **2** | **5** | **7** |
| Urumqi Blood Center |  | 2 | 5 | 7 |
| **Grand Total** | **113** | **60** | **107** | **280** |

**Figure 1.**
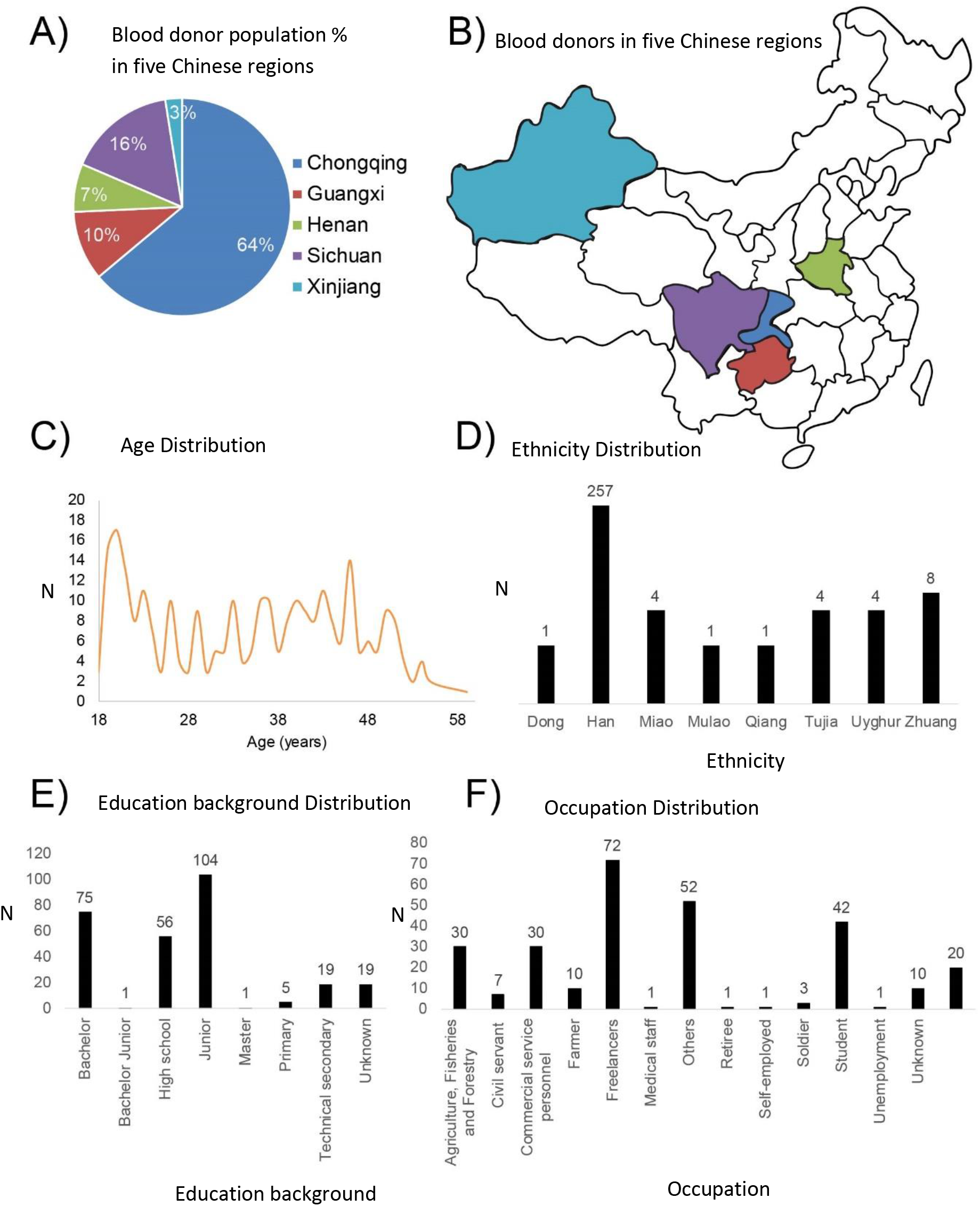
Demographic characteristics of blood donors

### HIV genotypes distribution

215 HIV samples were tested for nucleic acid, and all had detectable viral load. Out of 167 HIV with viral load >1000IU/mL, at least one sequence was obtained from 107 (64%) samples, not including 5’LTR. Some samples failed in sequencing may due to low viral load or RNA degradation during transport or sequence variations at the primer binding sites (26). Among 215 HIV positive samples,173 (80%) samples were collected in 2016, 22 (10%), and 20 (9%) were collected in 2015 and 2014, respectively. The majority of gender was male 177 (82%). About 99 (46%) HIV infected individuals’ education level was higher than or equivalent to a high school degree. 71 (33%) individuals were Junior degree. Similar results were described in a previous report (27). The freelancers (53, 25%) were major infected populations, followed by commercial service personnel (28, 13%), then farmer (14, 7%), and others (45, 20%) with the unknown occupation.

Two fragments, *env* IDR and *pol* INT, were taken into account for analysis except for the samples, which failed to obtain both of those two regions, to determine genotypes of 107 sequenced samples. Two genotypes B and C were identified with four types of CRFs (CRF 01, CRF07, CRF08, CRF55) and 5 URFs (URF_0107, URF_01C, URF_0708, URF_BU and URF_CU). The majority of samples genotype were C (42%), followed by CRF01_AE (24%) and CRF07 (17%) (Table 2), which was consistent with a previous report (28). In 45 genotype C samples, 15 samples, which successfully amplified both Env and Pol regions, were finally classified. There were 30 other samples classified in a single Pol region. There should be more C type recombinants if we can apply additional Env region for genotyping. Besides major types of C, CRF01_AE and CRF07, the residual CRFs and URFs were all other BC or BC and AE recombinant forms (data not shown)

**Table 2.**
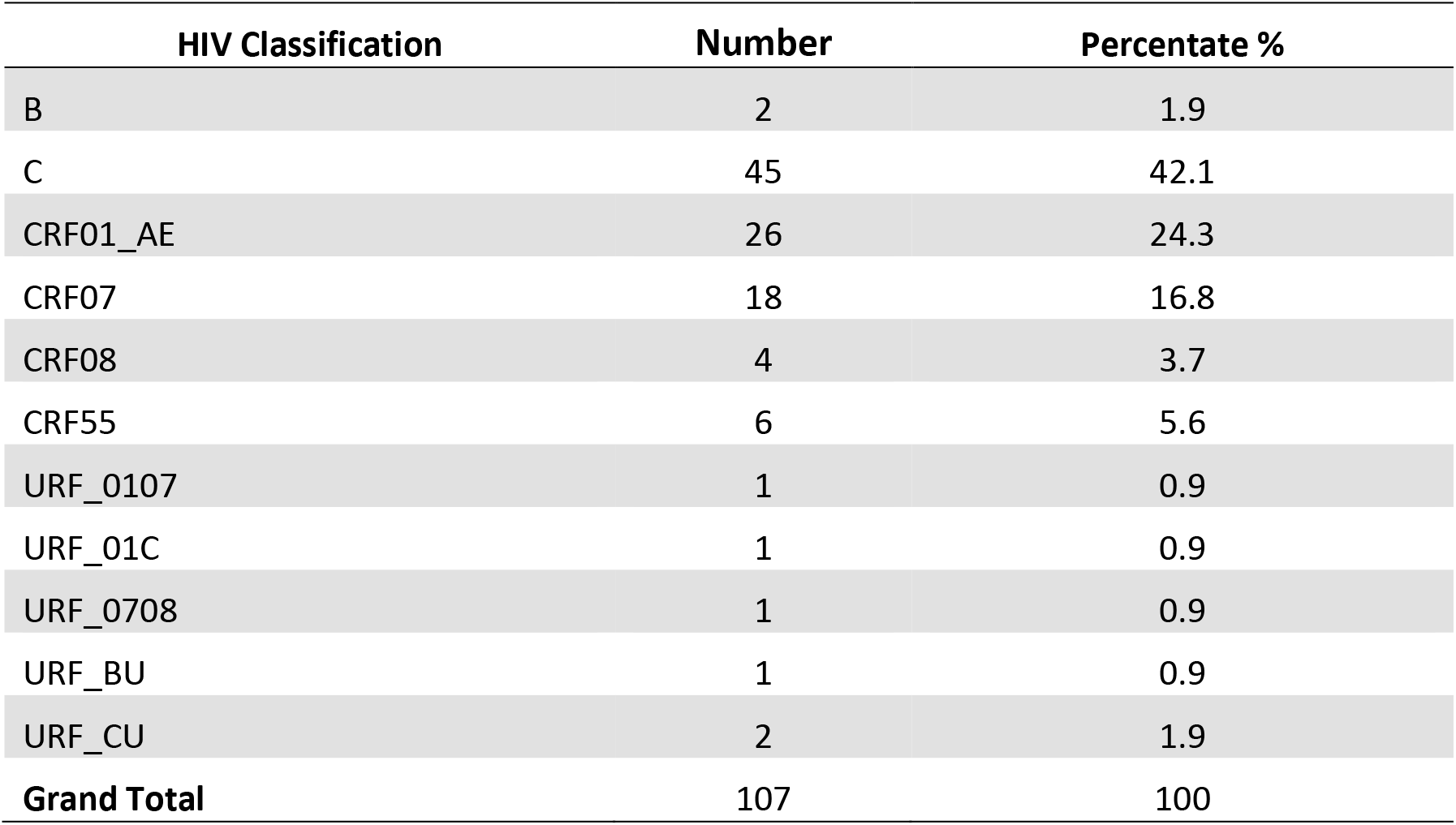
HIV classifications.

| HIV Classification | Number | Percentate % |
| --- | --- | --- |
| B | 2 | 1.9 |
| C | 45 | 42.1 |
| CRF01_AE | 26 | 24.3 |
| CRF07 | 18 | 16.8 |
| CRF08 | 4 | 3.7 |
| CRF55 | 6 | 5.6 |
| URF_0107 | 1 | 0.9 |
| URF_01C | 1 | 0.9 |
| URF_0708 | 1 | 0.9 |
| URF_BU | 1 | 0.9 |
| URF_CU | 2 | 1.9 |
| **Grand Total** | **107** | **100** |

Amongst 107 classified samples, 75% was from Chongqing and Sichuan, which is located in the Southwest of China (Table 3). The C was the majority genotype in Chongqing, Sichuan, and Xinjiang s. Genotype C, CRF01, CRF07 and CRF55 were almost equally distributed in Henan province. In Guang Xi province, the major genotype was CRF01, which was concordant with a previous study (28).

**Table 3.**
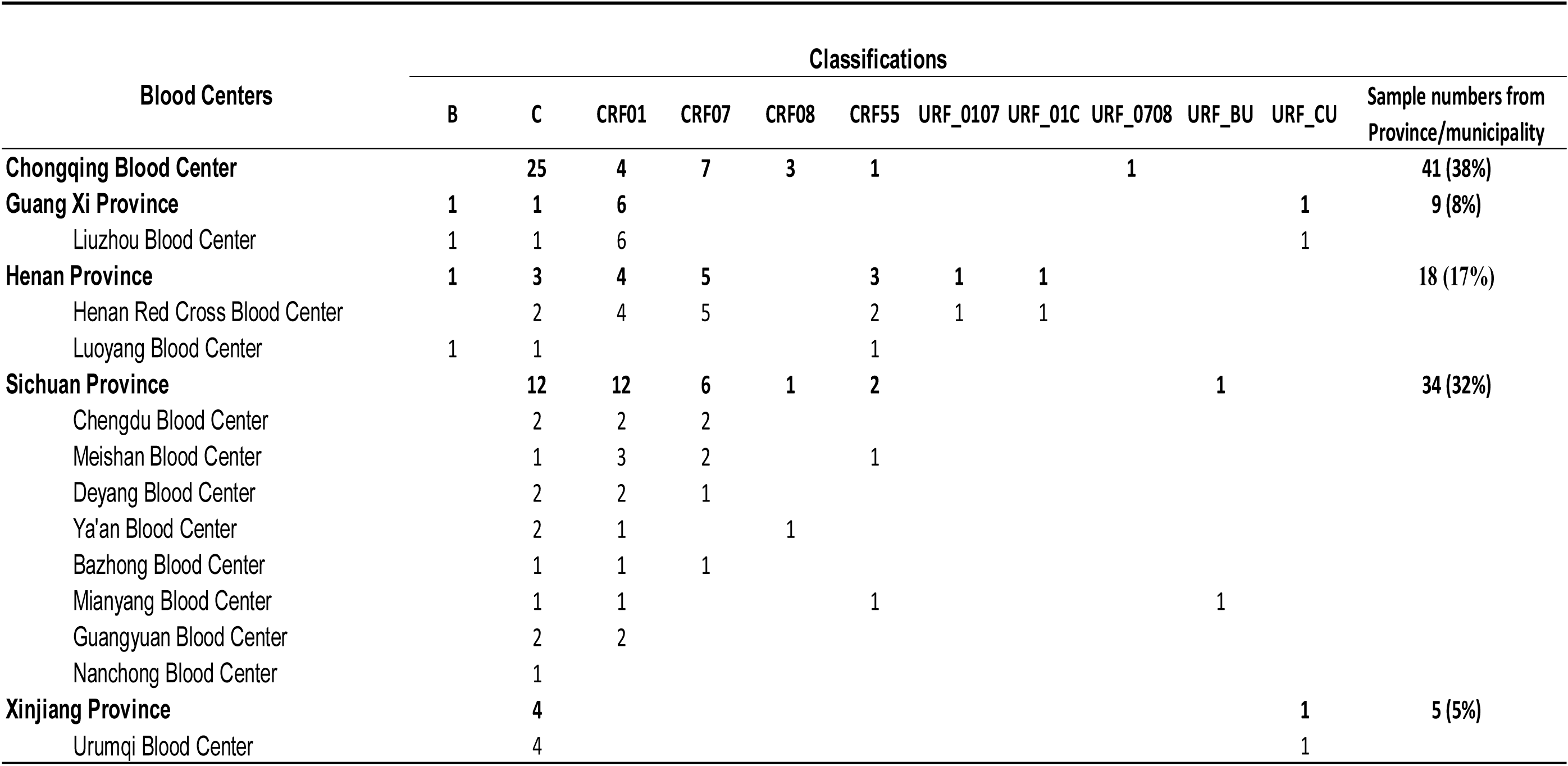
HIV distribution in different provinces

### HBV genotypes distribution and escape mutations

207 HBV samples tested for viral load were from Chongqing and Mianyang blood centers of Sichuan province located in the southwest of China. The mean donor age was 38.7± 11.3 years. The major ethnicity was Han (98.1%) with a few minorities of Dong, Miao and Tujia. The main education level was junior degree (37.4%), bachelor (31.2%) and high school degree (19.7%) (Table 1, E). A total of 120 samples were tested on M2000 and found VL> 100IU/ml. 113 among these samples had been successfully sequenced with at least one sequence. The HBV genotype/subgenotype was analyzed. The three genotypes B, C and D were identified (Table 4). Genotype B and C were predominant. Genotype B was the most common genotype (89.4%), and B2 was a major subgenotype (84.1%). 9.8% genotype C and 0.9% genotype D were identified. This was in line with a previous publication (29), but in our study, the proportion of genotype B was higher, which may be due to almost all samples analyzed being from a single Chongqing blood center located in the southwest of China.

**Table 4.** HBV classifications.

| HBV Classification | Number | Percentage % |
| --- | --- | --- |
| B | 6 | 5.3 |
| B2 | 95 | 84.1 |
| C | 1 | 0.9 |
| C1 | 2 | 1.8 |
| C2 | 8 | 7.1 |
| D | 1 | 0.9 |
| **Grand Total** | **113** |  |

The mutations in the S open reading frame region were analyzed as well. 79 out of 113 (69.9%) samples were found to have variants in “a” determinant region where antibodies predominantly target. The “a” determinant is a hydrophilic region of the HBV surface antigen (HBsAg) protein that was important for HBsAg detection by the immune response or diagnostic tests. 69 out of 79 (87.3%) samples had multiple mutations. And 19 (24.1%) had escape mutants (Figure 2), which can escape antibody detection and neutralization. The mutation rate was much higher than previous report 17.1% (29) in blood donors in China, also higher than reported elsewhere, including Japan (24%) (30), Korea (50%) (31), France (28%) (32) and Spain (40%) (33). In this study, there was no mutant identified in position 145, which was the most widely reported. The most frequent mutations found (Figure 2) in the study were located in position 133, then 129 and 134, which was different from a previous report (34).

**Figure 2:**
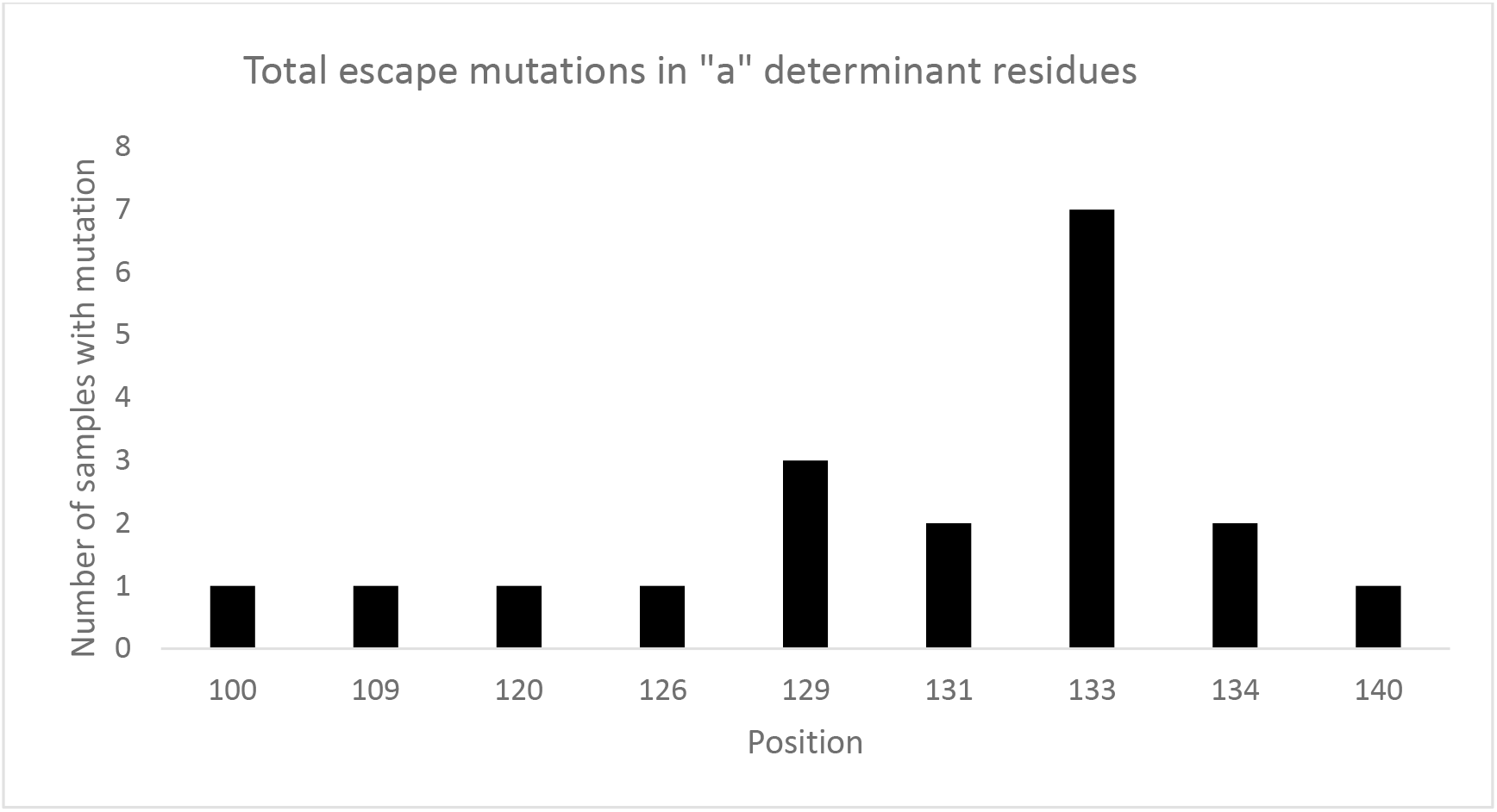
The distribution of escape mutants in “a” determinant region of HBV S region.

### HCV genotypes distribution

203 HCV samples were collected from 6 blood centers and were tested viral load positive, 60 (from 5 blood centers) were sequenced successfully with at least one sequence in the 5’UTR-Core region. 66.7% of donors were male and 33.3% were female. The mean age of donors was 36.2±10.6. The majority of ethnicity was Han (48, 80.0%) and with minorities Zhuang (6, 10%), Miao (1, 1.7%), Mulao (1, 1.7%), Qiang (1, 1.7%), Tujia (1, 1.7%) and Uyghur (2, 3.3%) (Table1, D). In total, 7 genotypes and subgenotypes were identified, namely, 6, 6a, 1a, 1b, 2a, 3a, 3b. The predominant genotype was 1b (N=29, 48%), followed by 6a (N=10, 17%), 2a (N=6, 10%), 3a (N=6, 10%) and small proportion of 3b, 6 and 1a (Table 5). The distribution of HCV classifications was concordant with a previous national wide investigation of HCV in mainland China, which reported the major HCV genotypes were 1b and 2a, accounting for 54.5% and 16.7%, respectively (35). In our study, 6a was a second major genotype. It was the dominant genotype in Liuzhou blood bank located in the south of China, which was similar to the publication of HCV subtypes of blood donors from 17 provinces and municipalities (36).

**Table 5.** HCV classifications.

| HCV Classification | Number | Percentage % |
| --- | --- | --- |
| 6 | 3 | 5.0 |
| 1a | 2 | 3.3 |
| 1b | 29 | 48.3 |
| 2a | 6 | 10.0 |
| 3a | 6 | 10.0 |
| 3b | 4 | 6.7 |
| 6a | 10 | 16.7 |
| **Grand Total** | **60** |  |

## Discussion

This is the first study to demonstrate that the strains of HIV, HBV and HCV commonly found in Chinese blood donors are detected by Abbott serology and molecular tests. Attempts to sequence and classify viremic samples are very important in providing evidence for blood screening safety in China.

HIV recombination is one of the main mechanisms contributing significantly to HIV-1 genetic diversity and unique recombinant forms are precursors of CRFs. Therefore, the monitoring of URFs will be helpful to the surveillance of new CRFs. CRF01_AE was found to be dominant in the transmitted sexual population in China (37), while CRF_07BC and CRF_08BC were mainly prevalent in IDU population (38). In addition, CRF07_BC was the most predominant strain in the north-western region, which suggested a lack of incoming transmissions from other regions (4). Although CRF01_AE only accounts for 5% of HIV cases worldwide, it plays an important role in China (39). In our study, the finding is in line with the previous study with the fact that the major CRFs are BC recombinant and the predominant CRF in the south of China is CRF01_AE. The limitation of the analysis is that most of the samples have one region (Pol or Env) involved in the analysis, which doesn’t have a breakpoint to judge whether it’s CRF07_BC or not. That’s why C is the majority genotype in our study. If we had complete genomes for classification, the CRF07_BC’s proportion should be higher.

Surveillance of HBV variants can help in monitoring the diversity and evolution of HBV. The investigation of HBV genotypes and mutations in blood donors will provide evidence to improve current sensitivity and specificity of screening assays and reduce TTIs in blood transfusion. According to the previous study (29), genotype B and C were the major genotypes in Chinese blood donors that is consistent with our study. However, other studies reported that genotype C has a higher prevalence in patients than genotype B (34, 40, 41), which is different from in blood donors. The most-reported mutant in “a” determinant is G145R which will affect HBsAg detection (21, 42). In our study, there is 69.9% of the HBV samples were found mutants in “a” determinant region with a distribution of escape mutants in "a" determinant region 69 (87.3%) multiple mutants and 19 (24.1%) escape mutants which will affect HBsAg detection. It indicates possible false-negative results of blood screening of HBsAg and further increase the potential threat of HBV to blood safety in China, which needs to be clarified by further studies.

Long-term multi-center molecular surveillance of HIV, HBV, and HCV amongst blood donors provides a detailed picture of the diversity distribution and variations, including new mutations and new subgenotypes occurrence in blood donors. It is very crucial to provide an efficient way to reduce transfusion-transmitted infections of HIV, HBV, and HCV and to improve blood safety further.

## Acknowledgements

The authors thank the technical support from Ana Vallari, Barb Harris, and Ana Olivo in the Abbott Global Surveillance Program. In addition, this work was supported by Abbott Diagnostics and the CAMS Innovation Fund for Medical Sciences (CIFMS, No.2016 - I2M - 1 - 018).

